# Deficiency of SYCP3-related XLR3 disrupts the initiation of meiotic sex chromosome inactivation in mouse spermatogenesis

**DOI:** 10.1101/2021.03.30.437712

**Authors:** Natali Sobel Naveh, Robert J. Foley, Katelyn R. DeNegre, Tristan C. Evans, Anne Czechanski, Laura G. Reinholdt, Michael J. O’Neill

## Abstract

In mammals, the X and Y chromosomes share only small regions of homology called pseudo-autosomal regions (PAR) where pairing and recombination in spermatocytes can occur. Consequently, the sex chromosomes remain largely unsynapsed during meiosis I and are sequestered in a nuclear compartment known as the XY body where they are transcriptionally silenced in a process called meiotic sex chromosome inactivation (MSCI). MSCI mirrors meiotic silencing of unpaired chromatin (MSUC), the sequestration and transcriptional repression of unpaired DNA observed widely in eukaryotes. MSCI is initiated by the assembly of the axial elements of the synaptonemal complex (SC) comprising the structural proteins SYCP2 and SYCP3 followed by the ordered recruitment of DNA Damage Response (DDR) factors to effect gene silencing. However, the precise mechanism of how unsynapsed chromatin is detected in meiocytes is poorly understood. The sex chromosomes in eutherian mammals harbor multiple clusters of *SYCP3*-like amplicons comprising the *Xlr* gene family, only a handful of which have been functionally studied. We used a shRNA-transgenic mouse model to create a deficiency in the testis-expressed multicopy *Xlr3* genes to investigate their role in spermatogenesis. Here we show that knockdown of *Xlr3* in mice leads to spermatogenic defects and a skewed sex ratio that can be traced to MSCI breakdown. Spermatocytes deficient in XLR3 form the XY body and the SC axial elements therein, but are compromised in their ability to recruit DDR components to the XY body.

**Author Summary:** A key event in the production of sperm is the pairing and synapsis of homologous chromosome pairs to facilitate genetic recombination during the first meiotic division. Chromosomal abnormalities that undermine pairing and synapsis in spermatocytes trigger a checkpoint that leads to removal of the abnormal cells via programmed cell death. In mammals, the sex chromosomes, X and Y, lack homology through most of their length and remain largely unpaired. To avoid triggering the checkpoint the X and Y are sequestered in a specialized nuclear compartment called the XY body. DNA damage response (DDR) proteins are recruited to the XY body in a highly ordered progression leading to repression of all gene transcription from the sex chromosomes. We show that knocking down expression of the X-linked *SYCR3*-like gene, *Xlr3,* disrupts sex chromosome gene silencing by interfering with recruitment of DDR factors, leading to compromised sperm production.

## Introduction

In eukaryotes, genetic recombination between paired, homologous chromosomes is enabled by induction of DNA double-strand breaks (DSBs) and repaired by factors of the DNA Damage Response (DDR) during meiosis I. Unpaired chromosomes or chromosomal segments are also subject to DSBs, yet they undergo DDR repair mechanisms specific to unsynapsed chromatin [1]. These repair mechanisms typically involve sequestration of unsynapsed chromatin in defined nuclear domains and subsequent transcriptional silencing and heterochromatinization in a process known as meiotic silencing of unpaired chromatin (MSUC) [2, 3].

In mammalian spermatogenesis, synapsis of the heterologous X and Y chromosomes is delayed compared to autosomes and is restricted to the pseudo-autosomal regions (PARs) [4, 5]. As a consequence, during the pachytene stage of prophase I a process consistent with MSUC occurs: a distinct nuclear compartment known as the XY body forms, wherein DSB repair depends on transcriptional inactivation of the sex chromosomes (MSCI) effected by DDR factors distinct from that of the synapsed autosomes [6]. MSCI initiates in pachynema and persists through meiosis and into spermiogenesis where the transcriptional repressed state is referred to as post-meiotic sex chromatin (PMSC) [7]. Most X and Y-linked mRNA-encoding genes are subject to silencing, although a few notable genes escape and are actively transcribed post-meiotically [8, 9]. Disruption of MSCI activates the pachytene meiotic checkpoint (PMC) and spermatocytes with active sex chromosome transcription undergo arrest and apoptosis [2].

The sensing of unsynapsed chromatin at the onset of homologous pairing is not well understood, but it appears to accompany the assembly of the synaptonemal complex (SC) and recruitment of the first DDR factors. After DSBs are induced, homologous chromosomes begin to pair and synapse as the SC assembles [10]. Homologs are arranged in chromosome loops anchored at the axial elements (AE) of the SC. The AE are constituted by Synaptonemal Complex Proteins 2 and 3 (SYCP2, SYCP3) [11, 12], which are necessary for the early events of DDR including recruitment of RPA, HORMAD1/2 and ATR to DSBs [13]. For reasons that are unclear, the DDR cascade in early pachynema is slightly different within the XY body: DSBs attract the damage sensors BRCA1 and TOPBP1, and ATR displaces from the axes into the loops [14–17] where it phosphorylates H2AX, leading to deposition of the repressive chromatin mark histone 3 lysine 9 trimethylation (H3K9me3) [18], sumoylation (SUMO1) [14] and ultimately to MSCI.

The precise role of SYCP3 in sensing unsynapsed chromatin is unknown. *SYCP3* is a highly conserved SC component found in the genomes of most metazoans, typically as a single copy autosomal gene [19]. However, in eutherian mammals numerous *SYCP3*-like amplicons are found across the X and Y chromosomes. The amplicons comprise the mammalian *X-linked Lymphocyte Regulated (Xlr)* superfamily [20] which includes: *SYCP3-like X-linked (Slx)* and *SYCP3-like Y-linked (Sly)* genes found in certain species of *Mus;* the *Xlr3, 4* and *5* triad found in a variety of Rodentia, and the *FAM9A, B* and *C* genes in primates [21–24]. The function of most *Xlr* members is unknown, however Cocquet and co-workers have shown that *Slx/Slxl1* and *Sly* are expressed in post-meiotic spermatids where they play antagonistic roles in the maintenance of PMSC [22, 25, 26]. Moreover, copy number variation of *Slx/Slxl1* and *Sly* between species of *Mus* is thought to underlie male sterility in inter-subspecific hybrids underscoring the importance of the *Xlr* family members in spermatogenesis [27].

Here we investigate the function of the *Xlr3* genes utilizing a transgenic mouse capable of tissue-specific expression of a short-hairpin RNA targeting *Xlr3* mRNA. Unlike other *Xlr* family members, which are typically present in dozens of copies, the *Xlr3*family consists of three closely-linked, protein-encoding paralogs, *Xlr3a, Xlr3b,* and *Xlr3c.* These genes are broadly expressed in mouse fetal and adult tissues and one paralog, *Xlr3b,* is imprinted in developing and adult mouse brain where it is expressed predominantly from the maternal X [28]. The three functional copies, hereafter referred to as *Xlr3,* encode a near-identical 26 kDa protein that is expressed in testis [21]. We show that *Xlr3* expression initiates during early meiotic prophase I where the protein localizes to the XY body in primary spermatocytes. Germ-cell specific knockdown of *Xlr3* mRNA in shRNA-transgenic males leads to partial disruption of spermatogenesis, reduced sperm count and offspring with a skewed sex ratio. Examination of sex-linked gene expression and localization of DDR factors points to a breakdown in the earliest stages of MSCI in the *Xlr3* knockdown males, implicating *Xlr3* as the earliest acting factor involved in XY body formation and the only known sex-linked factor implicated in MSCI.

## Results

### *Xlr3* is a multicopy gene encoding an *Sycp3-*like Cor1 domain

*Xlr* superfamily genes are scattered in multicopy clusters across the mouse X chromosome. The *Xlr3a/b/c* paralogs map to a small cluster encompassed within ~250 kilobases on mouse XA7.3 (Fig. 1A). *Xlr3a/b/c* encode near-identical proteins; XLR3B and XLR3C contain a single residue difference, while XLR3A contains eight amino acid substitutions compared to the other two paralogs (Fig. 1B). Our current understanding of structure/function relationships of *Xlr* superfamily members is informed by structural studies of the family progenitor, *Sycp3.* The N-terminus of SYCP3 binds DNA to facilitate the scaffolding of homologous chromosomes. Central coiled-coil helices are formed by the Cor1 domain, which tetramerizes to create SYCP3 fibers, while assembly of fibers into larger filaments is governed by the C-terminal region [29]. While the Cor1 domain is the most highly conserved segment among *Xlr* superfamily members, they all possess a monopartite nuclear localization signal (RKRK) within the divergent N-terminus (Fig. 1C).

**Figure 1.**
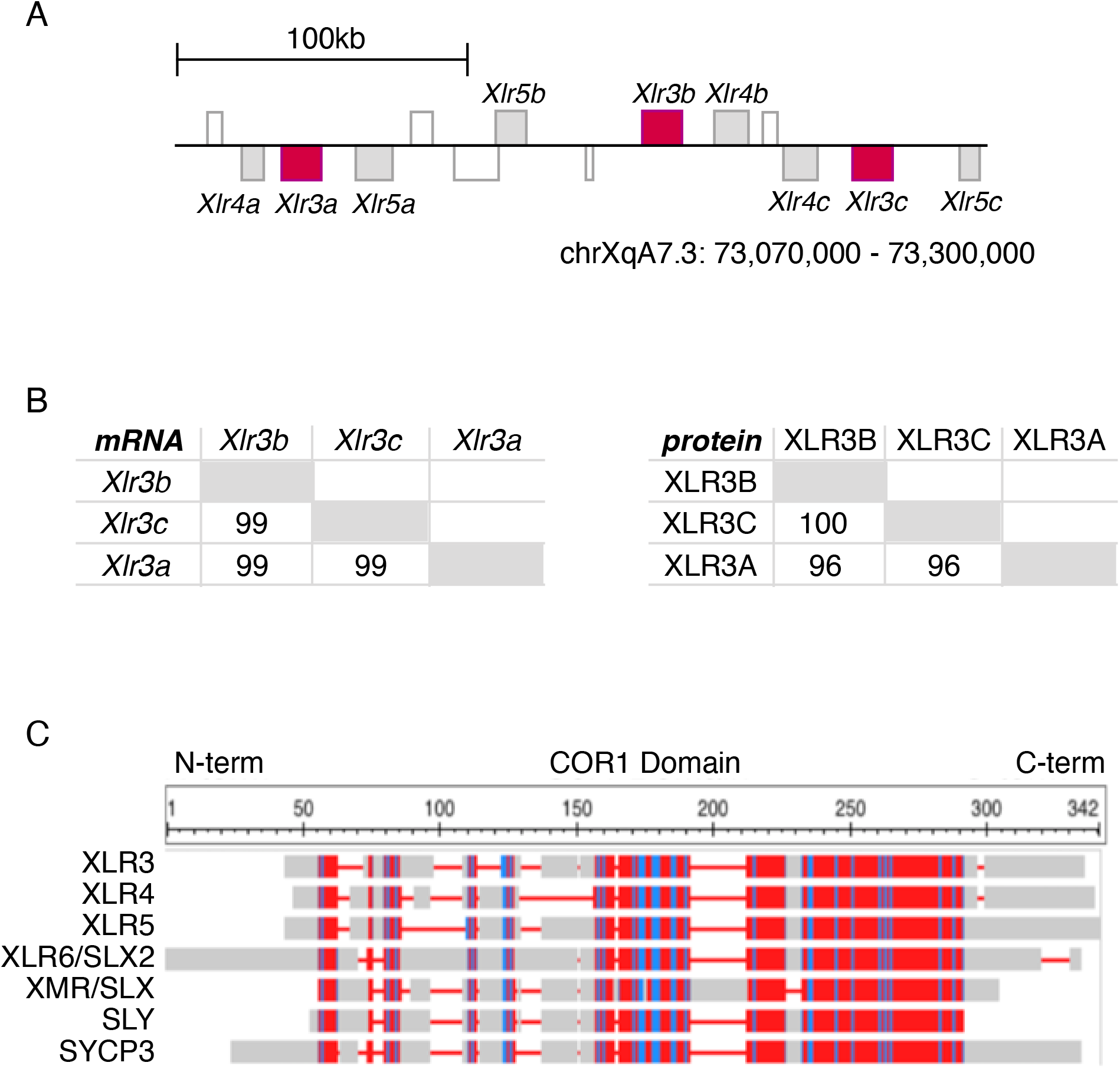
*Xlr3* is a family within the *Xlr* superfamily with X-linked copies and a Cor1 protein domain. **A)** The three open reading frame-containing functional *Xlr3* copies are located on *Mus musculus* X7A.3. Diagram based on mm10 genome build. **B)** Percent identity matrices of the functional *Xlr3* paralogs at the mRNA (left) and protein (right) levels generated using ClustalW. **C)** The protein structure encoded by the XLR A/B/C paralogs have a COR1 protein domain and similar domain organization compared to other XLR superfamily members from the N-to C-terminal regions. Color indicates residue conservation wherein gray indicates in/del sites, and red or blue indicate high or low side chain similarity, respectively. Diagram generated with amino acid sequences aligned by COBALT (NCBI).

### *Xlr3* is upregulated in early prophase I and colocalizes with the XY body in pachynema

To quantify *Xlr3* transcript levels during spermatogenesis, quantitative reverse transcription PCR (qRT-PCR) using primers common to the three *Xlr3* mRNA paralogs *(a/b/c)* was used. In the testes of staged-prepubertal mice, *Xlr3* transcription begins in pre-meiotic stages, but increases sharply from 8.5 days post-partum (dpp), representing meiotic entry, through 10.5dpp, representing prophase I leptonema (Fig. 2A). *Xlr3* transcript abundance peaks by 11.5dpp, at which point cells enter and progress through zygonema (Fig.2A).

**Figure 2.**
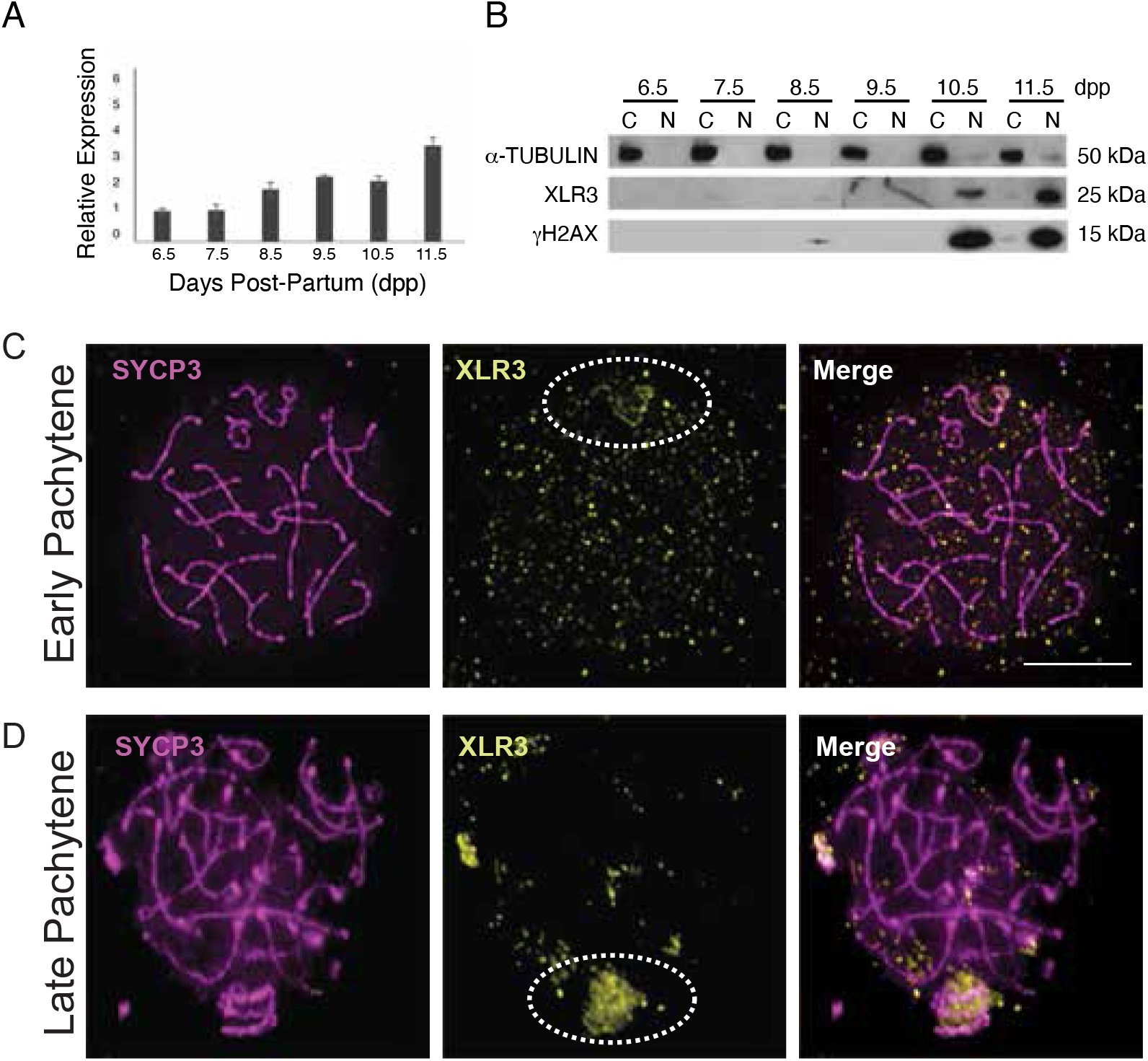
*Xlr3* expression is regulated stage-specifically in mouse spermatogenesis and localizes to the XY body during meiosis I. **A)** *Xlr3* mRNA expression increases during pre-leptotene stages (8.5-9.5 dpp) and is three-fold higher in zygotene (11.5 dpp) than 6.5 dpp. **B)** The XLR3 protein is only observed in the nuclear fraction from 10.5 dpp and strongly at 11.5 dpp through leptonema/zygonema. **C-D)** Immunocytochemistry of mouse spermatocyte chromosome spreads with DAPI (grey), SYCP3 (magenta), and XLR3 (yellow) reveals XLR3 localization. **C)** During early pachytene, XLR3 colocalizes with SYCP3 on the X and Y chromosome axes, indicated with a dotted circle. **D)** By late pachytene, XLR3 moves to form a cloud around the XY bivalent, indicated with a dotted circle. Scale bar = 6 *μ*m.

While *Xlr3* transcripts are detected before meiotic entry, XLR3 protein is not detectable via immunoblot until 10.5-11.5dpp, where it appears restricted to the nucleus (Fig.2B). Thus, XLR3 translation and nuclear localization coincide with the appearance of DSBs during leptonema [6]. Through immunocytochemistry (ICC), we observed specific XLR3 protein subcellular localization to the XY body during pachynema (Fig.2C-D). In early pachynema, XLR3 closely associates with the sex chromosome axes, including on the PAR (Fig.2C). By late pachynema, XLR3 appears to move away from the axes, but maintains association with the XY body, suggesting it translocates to the XY chromatin loops (Fig.2D).

### shRNA*-Xlr3* activity significantly reduces *Xlr3* abundance in mouse testis

Upregulation in early prophase I and localization to the XY body suggests a possible role for *Xlr3* in sex chromosome regulation in meiosis I. To explore this possibility, we created a mouse model in which *Xlr3* function is abrogated through RNA interference (RNAi) in an approach similar to that described by Cocquet *et al.* [22, 25]. To knock down *Xlr3* post-transcriptionally, we designed a construct containing a short hairpin RNA (shRNA) with sequence complementary to the *Xlr3* exon3/4 region (Fig. 3A) that is invariant among the *Xlr3* paralogs but divergent from *Xlr4* and *Xlr5* at several sites (Fig. S2A). A floxed stop cassette between the *ROSA26* promoter and the *Xlr3*-shRNA transgene was excised by crossing to a *Ddx4-Vasa* Cre recombinase mouse, allowing initiation of expression of the shRNA in spermatogonia [30] (Fig. S1). Efficacy of the shRNA knockdown was assessed via qRT-PCR using primers targeting exon 3 of *Xlr3,* the site of shRNA binding, and primers further downstream at exon 6. *Xlr3* transcript abundance was decreased by approximately 50% in shRNA-*Xlr3* mouse testis compared to that of age-matched wild type males (Fig. 3B). Immunofluorescence (IF) was used to quantify the effect of this knock down on XLR3 protein levels in spermatocytes. Using γH2AX as a marker for the XY body, we quantified the XLR3 signal overlap in wild type and shRNA-*Xlr3* pachytene spermatocytes. Compared to the wild type, shRNA-*Xlr3* cells had on average approximately 50% of the IF signal (Fig. 3C), suggesting the reduction of protein product is directly proportional to that of *Xlr3* transcript levels.

**Figure 3.**
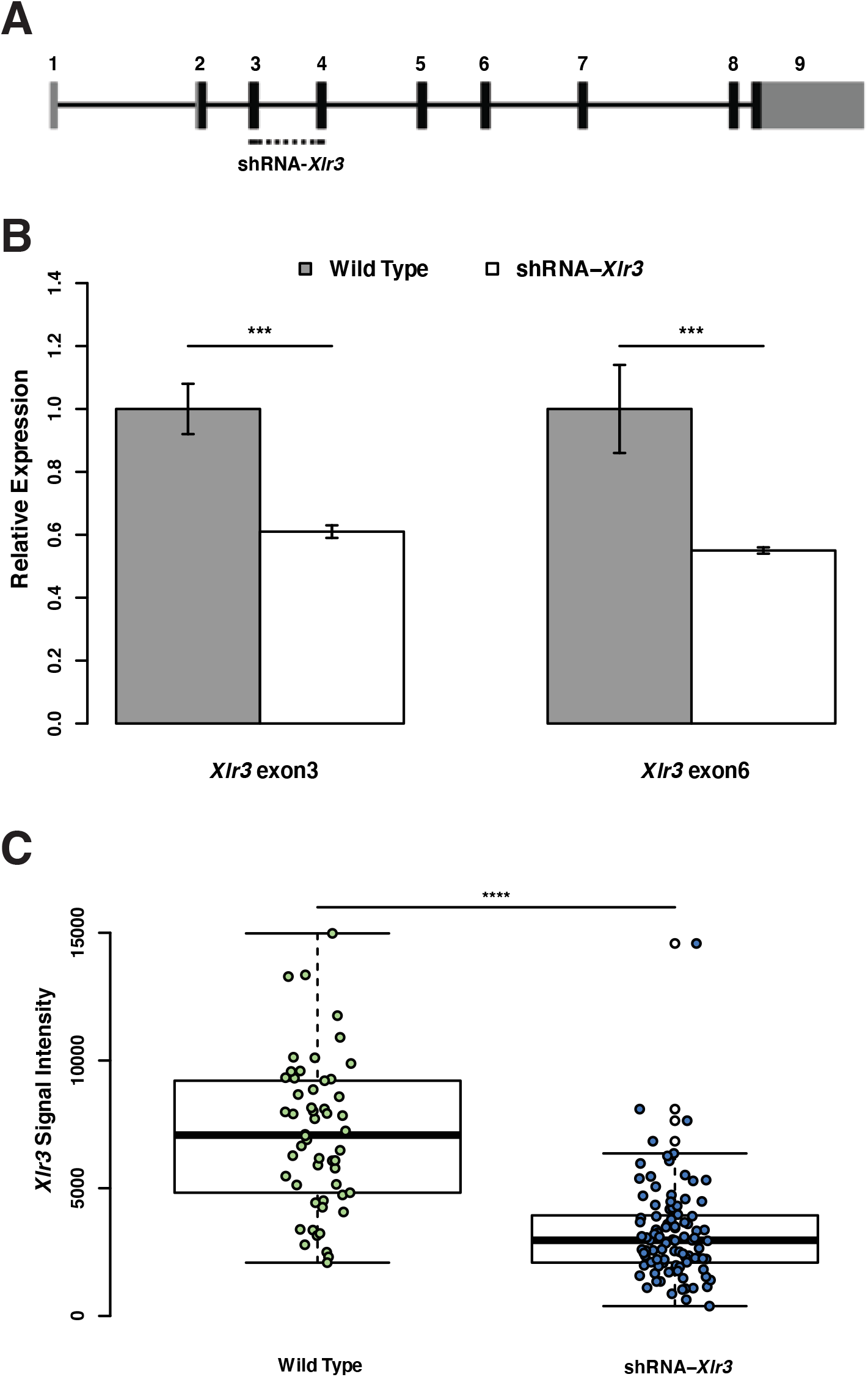
The shRNA-*Xlr3* targets transcripts at exon 3 and transgenic mice display *Xlr3* knock down. **A)** Diagram of *Xlr3* gene sequence and the location of shRNA-*Xlr3* targeting. Gray and black boxes represent UTRs and ORF regions, respectively. **B)** *Xlr3* mRNA is knocked down ~50% at 9.5 dpp in shRNA-*Xlr3* compared to age-matched wild type mice. Expression levels were assessed by qRT-PCR of whole testis cDNA. Two primers were used to measure the transcript at the location of shRNA targeting, exon 3 (Paired t-test, df=5, p-value=0.0001016***), and downstream, exon 6 (Paired t-test, df=5, p-value=2.725e-05***). Error bars represent mean C.I. **C)** XLR3 signal on the XY body is reduced in 17.5dpp spermatocytes. Protein levels were assessed by immunofluorescence of surface spread spermatocytes from shRNA-*Xlr3* and age-matched wild type cells (Wilcox Test: W=646, p-value=1.019e-14****).

To verify specificity of the shRNA to *Xlr3,* we used qRT-PCR to assay mRNA levels of the most closely related *Xlr* family members, *Xlr4* and *Xlr5.* Transcript levels of these genes were unaffected in the testis of shRNA-*Xlr3* transgenic mice (Fig. S2B,C). To assess potential induction of a viral immune response due to the expression of double-stranded RNA in the transgenic mice, we assayed expression of *2’,5’-Oligoadenylate synthetase 1b (Oas1b),* which is part of the interferon viral response pathway [22, 31, 32]. Again, there was no significant difference in transcript level of this gene, indicating our shRNA did not induce an interferon response (Fig. S2D). Taken together, these data indicate we have achieved specificity of the targeted shRNA to *Xlr3* reduction in testes.

### *Xlr3* knock down leads to germ cell loss and a skewed sex ratio

Initially, both male and female shRNA-*Xlr3* transgenic mice appeared to be fully fertile, as they produced offspring with the same frequency and litter size as wild type sibling controls (Fig. 4A-B). However, a thorough examination of the transgenic males revealed a significant reduction in the sperm count (49.3% reduction) and normalized testes weight (14.7% reduction) at 14 weeks of age compared to wild type siblings (Fig. 4C-D). Histological cross-sections of the testes of shRNA-*Xlr3* transgenic males revealed the presence of disorganized seminiferous tubules and a significant decrease in average tubule diameter (50% reduction) suggestive of either germ cell or Sertoli cell loss (Fig. 4E-G). To distinguish between these two possible causes of tubule defects, we assayed expression of the Sertoli cell marker, *Sox9* (Rebourcet et al., 2014, 2017) to test for Sertoli cell loss. There was no difference in *Sox9* expression between transgenic testis and wild type, suggesting Sertoli cell loss has not occurred (Fig. S2E). However, a twofold increase in germ cell apoptotic figures was observed in tubules of shRNA-*Xlr3* transgenic males compared to wild type (Fig. 4H-J). Overall, the *Xlr3* knock down leads to significant germ cell loss, but not to an extent to subvert fertility.

**Figure 4.**
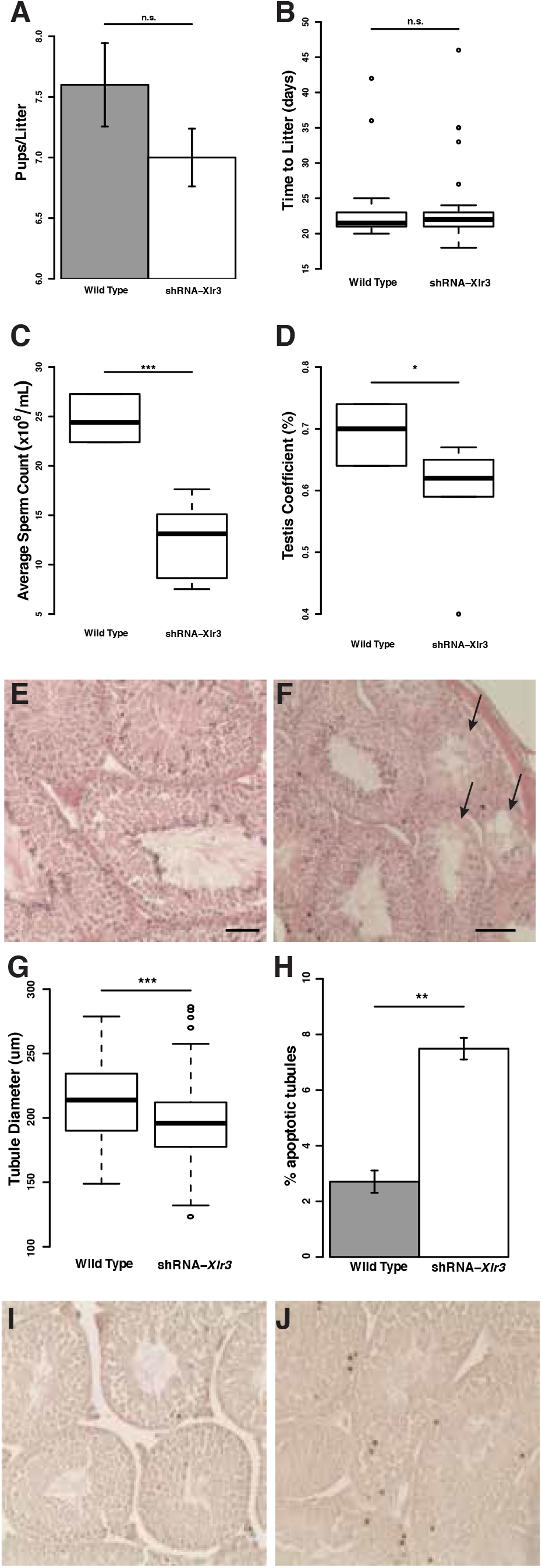
*Xlr3* knock down mice are fertile but have spermatogenic defects. **A)** Across 50 shRNA-*Xlr3* sired litters, the number of pups produced per litter is not significantly (n.s.) different from that of wild type counterparts (Wilcox Test, W=135, p-value=0.7778) and **B)** there is no significant difference in the time to litter (T-test, t=0.2017, df=52.434, p-value=0.8409). **C)** Upon further examination of a cohort of 14-week old males, significantly reduced sperm count (Paired t-test, df=6.1897, p-value=0.0005***) was observed, as well as **D)** significantly reduced testis weight as percent of body weight (Paired t-test, df=6.8775, p-value=0.03966*). **E,F)** H&E staining revealed that shRNA-*Xlr3* testes **(F)** have several disorganized seminiferous tubules, indicated by arrows, that are not present in any wild type section **(E)**. **G)** The tubules of these mice are also significantly smaller in diameter than those of their wild type counterparts (T-test, df=9694, p-value=6.889e-06****). **H-J)** These phenotypes may be due to **(H)** an increased number of apoptotic cells per tubule (T-test, df=3, p-value=0.006456**), which was observed through TUNEL stained wild type **(I)** and shRNA-*Xlr3* **(J)** testes. Apoptotic cells are stained dark brown. Error bars represent mean ± C.I. Scale bars = 20 *μ*m.

During the course of fertility assessment, we observed a significant skew (60:40) towards female offspring produced by the shRNA-*Xlr3* males across 400 individuals (Fig. 5A). To determine if sex chromosome nondisjunction (ND) events in the transgenic males were leading to excess daughters (i.e. X^mat^ monosomics), we mated shRNA-*Xlr3* males on a C57BL/6J background to wild type C3H/HeJ females to trace the parental origin of the X chromosome via strain-specific sequence polymorphisms (Fig. 5B). While the same 60:40 skew toward females was observed in the inter-strain cross, only 2% of hybrid offspring were found to be 39,X^mat^, attributable to sex chromosome ND (Fig. 5C). While this rate of ND is significantly higher compared to spontaneous events (0.01%) [33], ND does not fully account for the observed increase in female offspring, indicating the *Xlr3* knockdown enhances X sperm or compromises Y sperm function.

**Figure 5.**
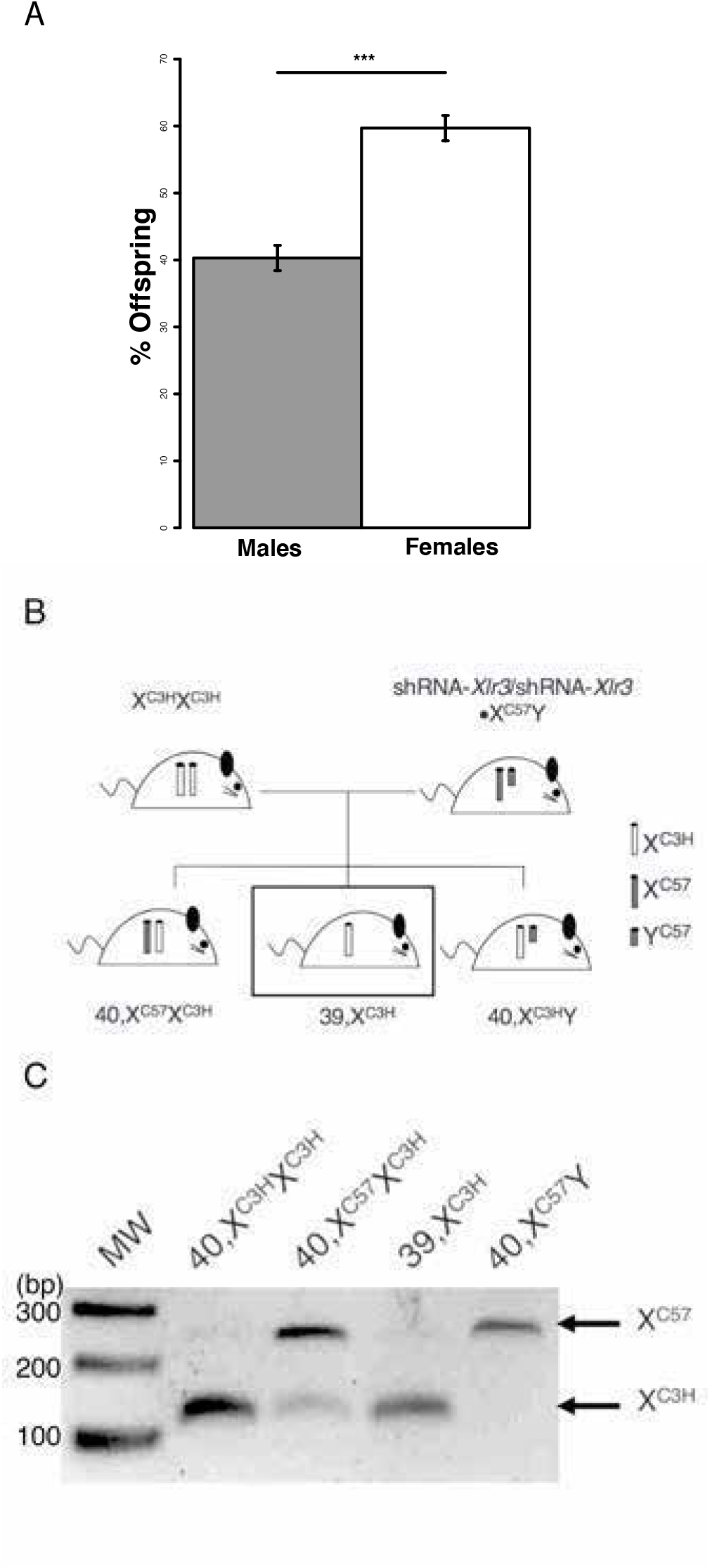
*Xlr3* knock down alters offspring sex ratios and increases XY non-disjunction. **A)** shRNA-*Xlr3* males produce a significantly increased percentage of female offspring than expected (400 offspring, binomial distribution test, p=0.5, p-value=0.0003238) compared to the expected 50:50 sex ratio. **B)** To identify XY nondisjunction, shRNA-*Xlr3/*shRNA-*Xlr3*•C57 males were mated to wild type C3H females to use polymorphism genotyping. We observed the production of ~2% hybrid 39,X offspring (box). **C)** ND events (X monosomy) were identified by PCR and *MseI* restriction digest. Ladder (MW) is shown to the left and X-specific allele is indicated to the right.

### *Xlr3* knock down leads to disruption of meiotic sex chromosome inactivation

RNAi knockdown of the *Xlr* superfamily members *Slx* and *Sly* also leads to sex chromosome transmission distortion and either enhancement (*Slx* knockdown) or disruption (*Sly* knockdown) of normal transcriptional repression of post-meiotic sex chromatin (PMSC) [22, 25]. Given the localization of XLR3 to the XY body in pachynema, we examined the status of MSCI in shRNA-*Xlr3* mice. To quantify changes in relative abundance of sex-linked transcripts, we assayed several genes by qRT-PCR at two time points. Since steady-state transcript levels are reflective of both transcriptional output and mRNA half-life, we determined potential changes in transcript levels in transgenics in reference to a normalized ratio of wild type transcript levels from 9.5dpp, the day before XLR3 protein is detected, and 14.5dpp, (Fig. 2B) the earliest point at which MSCI is detected [34]. We observed a statistically significant upregulation of several genes across the X chromosome *(Rhox13, Hprt, Fmr1, Rps6ka6, Tcp11×2),* as well as the “pachytene-lethal” Y-linked gene *Zfy2* [10, 35] (Fig. 6A-B).

**Figure 6.**
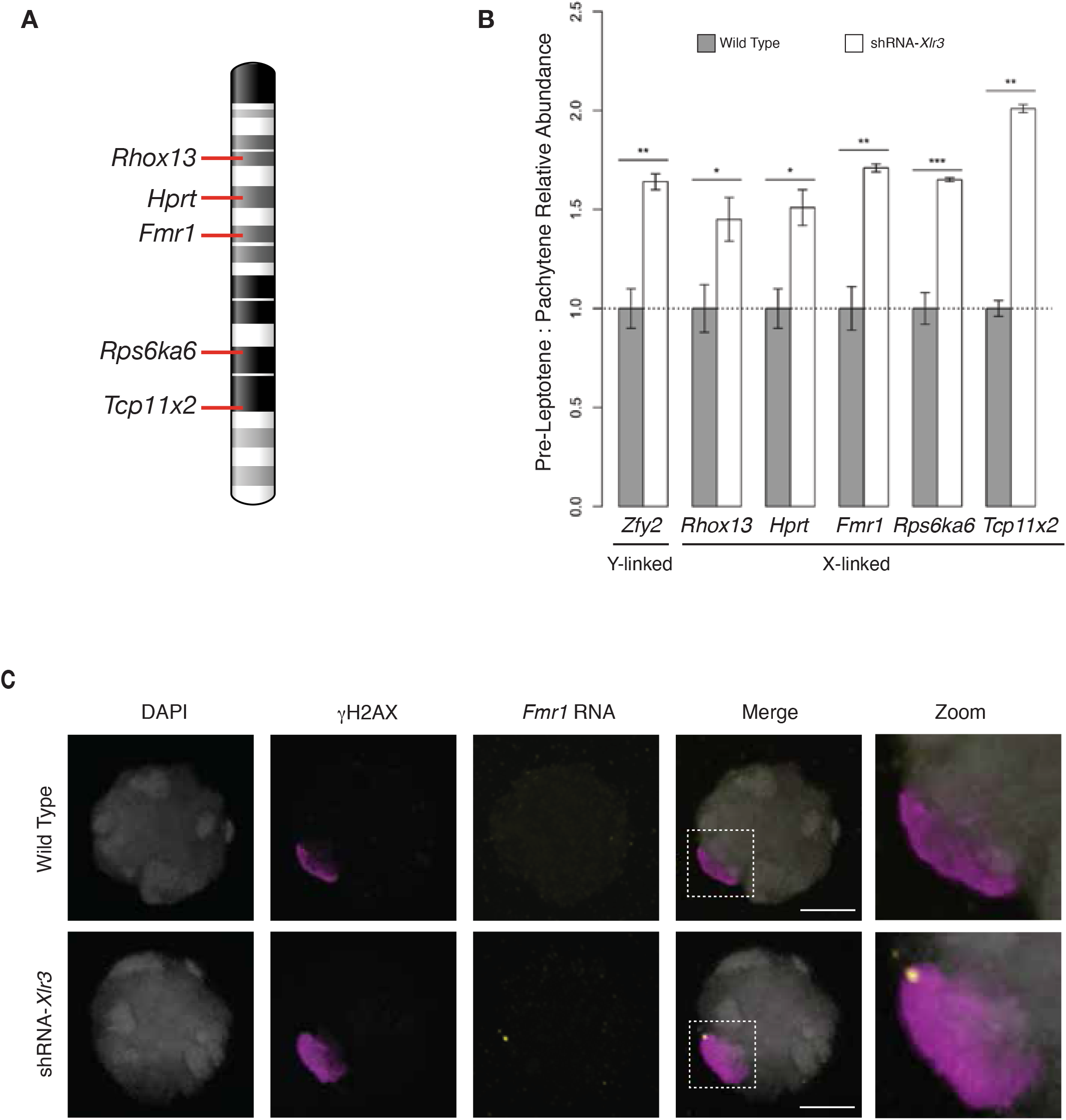
*Xlr3* knock down disrupts MSCI. **A)** Map (ideogram of mm10) of X-linked targets. **B)** To detect transcript abundance of the X-linked targets and *Zfy2* during pachytene (P) when the XY body should be silent, transcript levels from 9.5 dpp (leptotene, N=3) and 14.5 dpp (pachytene, N=3) testes were measured by qRT-PCR and normalized to those of age-matched wild type mice. All targets were significantly upregulated by the knock down of *Xlr3* (One-sided T-test, *Zfy2:* t=-8.3929, df=2.4548, p-value=0.003624, *Rhox13,* t=-2.8488, df=2.1466, p-value=0.04812, *Hprt,* t=-4.2463, df=2.0958, p-value=0.02358, *Fmr1,* t=-40.279, df=2.9412, p-value=1.984e-05, *Rps6ka6:* t=-13.992, df=2.8365, p-value=0.0005269, *Tcp11×2:* t= −5.8356, df=2.5986, p-value=0.007383). **C)** RNA FISH of 17.5dpp spermatocytes using *Fmr1* probe (yellow), γH2Ax as an XY body marker (magenta) and DAPI as a counterstain (grey). No overlap of *Fmr1* and γH2Ax was observed in the wild type cells (top), while signal overlap was observed in the shRNA-*Xlr3* transgenic cells (bottom). Far right panel is zoom of boxed inset shown in merge images. Scale bars = 6 *μ*m.

Since qRT-PCR measures abundance of transcripts, but cannot differentiate between stable transcripts and those actively transcribed at any one time point, we used RNA fluorescence *in situ* hybridization (RNA-FISH) to verify active transcription from the XY body during pachynema. Using a probe for *Fmr1,* one of the X-linked qRT-PCR targets (Fig. 6A-B), we confirmed there are no nascent transcripts observable within the XY body (0/40 cells) in wild type spermatocytes that maintain active MSCI regulation (Fig. 6C). However, approximately 50% of spermatocytes from shRNA-*Xlr3* homozygotes showed colocalization of the *Fmr1* signal with the γH2AX XY body marker (29/60 cells), indicative of MSCI disruption commensurate with the extent of *Xlr3* knock down (Fig. 6C).

### XLR3 marks the XY body to recruit DDR and chromatin regulators

Since XLR3 production and nuclear localization coincides with the appearance of DSBs in meiotic prophase I, it is important to place XLR3 localization to the XY body in relation to the sequential recruitment of DDR factors that bring about MSCI. BRCA1 recruitment to sex chromosome DSBs in late zygonema/early pachynema leads to localization of ATR kinase to sex chromosome axes (Turner et al., 2004). Subsequently, ATR translocates from the axes to chromatin loops in early to mid-pachynema where it phosphorylates H2AX [16, 17, 36], and is followed by the later accumulation of SUMO1 [37, 38], and H3K9me3 [18] by diplonema.

We assessed the sequential localization of XY body markers via ICC from later to earlier stages (from top to bottom Fig. 8). In late-pachytene spermatocytes (18.5 dpp) of the *shRNA-Xlr3* males we detected a spectrum of signal intensity for SUMO1 (Fig.7A-C). While some spermatocytes showed staining equivalent to wild type, many in the same spreads showed severely diminished staining. The average corrected total fluorescence (CTF) of the anti-SUMO1 signal was half the intensity in shRNA-*Xlr3* cells compared that of WT cells (Fig. 7A-C). Contrarily, the accumulation of H3K9me3 at the XY body, another late pachytene signal, was found to be elevated on average in shRNA-*Xlr3* cells compared to wild type, again showing a spectrum of signal (Fig. 7D-F). These results suggest that XLR3 plays a role in regulating the epigenetic landscape of the XY body, resulting in higher levels of H3K9me3 at the XY body at the end of pachynema [18].

**Figure 7.**
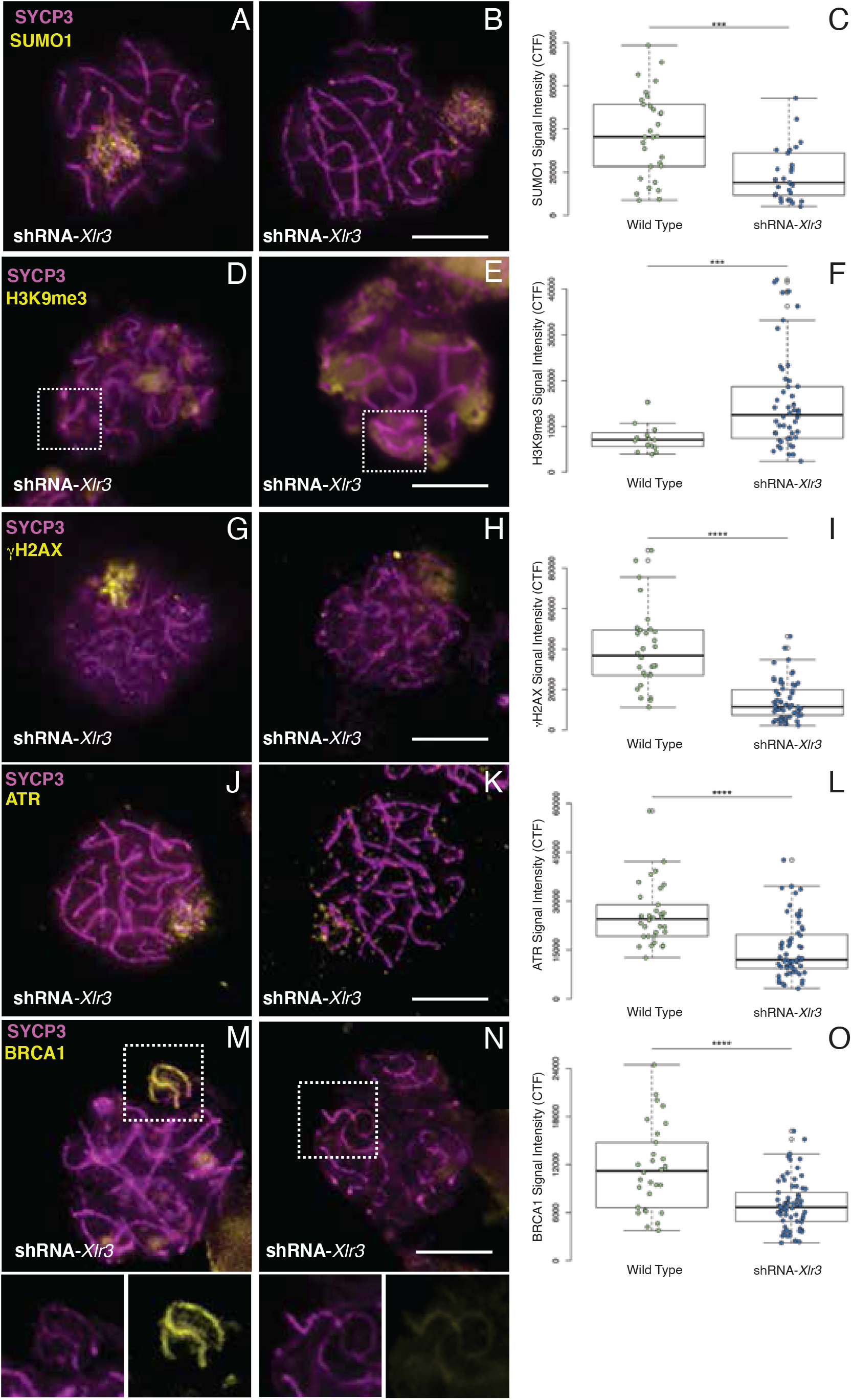
XY body MSCI DDR factors and chromatin modifications are diminished in shRNA-*Xlr3* pachytene spermatocytes. Surface spread images were subjected to ICC with an antibody against SYCP3 (magenta). **A-E)** DDR factor signal intensity (yellow) in shRNA-*Xlr3* spermatocytes is equal to that observed in wild type cells (left) and is diminished in some knockdown cells (middle). Overall, the average shRNA-*Xlr3* population DDR signal corrected total fluorscence (CTF) were significantly decreased (right). **A-C)** SUMO1 signal across the XY body loops (Wilcox Test: W = 188, p-value = 0.000497***). **D-F)** H3K9me3 signal across the XY body loops (T test: t = 4.6677, df = 62.88, p-value = 1.644e-5***), **G-I)** γH2Ax signal across the XY body loops (Wilcox Test: W = 169, p-value = 5.298e-10****), **J-L)** ATR signal across the XY body axes and loops (Wilcox Test: W = 384, p-value = 5.713e-07****), **M-O)** BRCA1 signal across the XY body axes (Wilcox Test: W = 1575, p-value = 3.995e-05****). Images are deconvolved quick projections. Scale bars = 10 μm.

In mid-pachytene spermatocytes (16.5 dpp), the average signal intensity for γH2AX at the XY body in shRNA-*Xlr3* transgenic mice is half that of wild type, again showing cell-to-cell differences from normal to weak staining (Fig. 7G-I). Likewise, ATR was observed to localize appropriately to the XY body, i.e. along the chromosomal axes and chromatin loops, but at approximately 50% average intensity in shRNA-*Xlr3* compared to wild type cells (Fig. J-L). Finally, BRCA1, the earliest known mark on the XY body preceding recruitment of ATR to the XY body [17], localized to the chromosomal axes of the XY body in early pachytene spermatocytes (13.5 dpp) in both transgenic and WT testis, but at half the level in shRNA-*Xlr3* transgenic cells compared to WT cells, again showing cell-to-cell differences in the knockdown (Fig. M-O). These results suggest that XLR3 localization is a necessary pre-condition for the recruitment of BRCA1 to the XY body, placing XLR3 as one of the earliest known factors leading to the demarcation of the XY body and the only known sex-linked factor essential in MSCI.

## Discussion

The first identification of an *X-linked lymphocyte regulated (Xlr)* gene came from the isolation of a B lymphocyte expressed cDNA that was an early candidate for a mouse X-linked immunodeficiency syndrome *(xid)* [39]. *Xlr* (a.k.a. pM1) was ultimately dismissed as *xid,* but deeper genomic characterization of this multicopy gene led to the discovery of other *Xlr* homologs with broad expression profiles scattered across the murine X and Y [21, 40, 41]. The *Xlr* name, with a few exceptions, has largely persisted despite the recognition that *SYCP3* is the ancestral source gene of this family [42, 43].

Sex-linked *SYCP3*-like genes can be found in the published genome sequences of a variety of eutherian mammals (e.g. *Canis lupus familiaris LOC102152531).* The broad distribution of these genes across Eutheria raises the question: Why have duplicate copies of *SYCP3* moved to the sex chromosomes where they have multiplied and diversified in the course of eutherian divergence?

At least some members of the *Xlr* family seem to have evolved under the influence of sexual antagonism as suggested by functional studies of the *Slx/Slxl1* and *Sly* genes [22, 25, 26, 44]. *Slx/Slxl1* and *Sly*, X-linked and Y-linked respectively, are multicopy genes found in certain species of *Mus* that are expressed predominantly in post-meiotic spermatids [42, 45]. Using a shRNA transgenic approach, upon which ours was modeled, Cocquet and coworkers [22, 25] showed knockdown of *Sly* results in subfertility, a female-skewed sex ratio, and relaxation of PMSC leading to up-regulation of X-linked genes. *Slx/Slxl1* knockdown males are also subfertile, but with the opposite effect observed in the *Sly* knockdown: PMSC repression is enhanced when *Slx/Slxl1* expression is reduced, leading to inappropriate silencing of several otherwise active X-linked genes and an offspring sex ratio skewed toward males. Curiously, full fertility is restored in double knockdown males expressing both anti-*Slx/Slxl1* and anti-*Sly* shRNAs [25], indicating that Y-linked copies are acting in conflict with X-linked copies [26]. Good and coworkers demonstrated that sterility in F1 hybrid males from the mating of two sub-species of *Mus musculus* is associated with a breakdown in PMSC, showing broad up-regulation of sex-linked genes [46]. The two sub-species differ significantly in copy number of the *Slx/Sly* genes; in hybrid males, the presence of a higher *Slx* copy number compared to *Sly* leads to the up-regulation of sex-linked genes, which is consistent with the induced *Slx/Sly* imbalance and PMSC failure in the knockdown model [25]. Recently, it was shown that SLX/SLXL1 compete with SLY in binding to the H3K4me3-reader, SSTY1, in spermatid nuclei, mediating their respective activities as transcriptional up-or down-regulators [26].

Unlike the *Slx/Slxl1/Sly* clusters, which are restricted to the genus *Mus*, the *Xlr3/4/5* cluster appears to be more ancient, having homologs with shared local synteny in dog, pig and alpaca. The broad distribution of these genes across eutherian lineages and our observations that knockdown of *Xlr3* results in disruption of BRCA1 and ATR localization to the XY body suggest a fundamental role for these genes in the initiation of MSCI in eutherian male meiosis.

Marsupials share a common origin for the sex chromosomes with Eutheria [47], and likewise share many features of MSCI in spermatogenesis, including formation of an XY body and localization of BRCA1 and ATR to the sex chromosomes early in meiotic prophase I [48, 49]. Unlike eutherians, the marsupial X and Y chromosomes lack a PAR and do not pair during meiosis I; no synaptonemal complex is formed but the sex chromosomes are held together within the XY body by an SYCP3-enriched structure called the “dense plate” [50]. The emergence of *Xlr* genes at the base of the eutherian lineage, therefore, suggests they were recruited into an established MSCI mechanism present in the therian common ancestor.

We envision two possible scenarios underlying the co-optation of a sex-linked *SYCP3* duplicate, *Xlr*, into MSCI. Like *Slx/Sly,* the earliest *Xlŕs* could have evolved as a consequence of the persistent intragenomic conflict between the sex chromosomes [51, 52]. Supporting this scenario is the observation that knockdown of *Xlr3* results in an offspring sex ratio skewed toward females (Fig. 5A). However, it is difficult to reconcile this result with the observation that knockdown of the only other X-linked *Xlr* members with known function, *Slx/Slxl1,* skews toward male offspring. In other words, normal *Slx/Slxl1* function appears to drives X chromosome transmission, while normal *Sly* function drives Y transmission. *Xlr3* appears to function more similarly to the Y-linked *Sly,* by promoting sex chromosome transcriptional silencing and, ostensibly, Y-chromosome transmission. While it is possible that *Xlr3* evolved as a suppressor of X meiotic drive, it is more likely that the sex ratio skew resulting from *Xlr3* knockdown is a byproduct of disrupted MSCI.

Alternatively, *Xlr* genes may have evolved to augment or adapt an ancestral therian MSCI mechanism to address an emergent feature of the sex chromosomes in Eutheria. The emergence of *Xlr* genes at the base of the eutherian lineage coincides with the addition of a large segment of autosomal DNA to the sex chromosomes after the divergence of Eutheria from Marsupialia [53, 54]. While this added segment likely paired initially, it is clear that the Y-borne segment underwent rapid attrition during the divergence of placental mammals [55]. We propose that *Xlr3* evolved to accommodate silencing of the emergent unpaired sex chromosome DNA. The factors that govern meiotic silencing of unpaired chromatin (MSUC) are drawn from a limited pool and, hence, are highly dosage sensitive [56–58]. If unpaired chromatin exceeds a certain threshold, transcriptional silencing is disrupted, leading to meiocyte loss or the production of aneuploid gametes [14, 59]. Likewise, it is clear that, like SLX/SLXL1/SLY in PMSC, XLR3 involvement in MSCI is dosage sensitive. Our finding that ~50% knockdown of *Xlr3* results in ~50% loss of sperm suggests that the compromised expression of *Xlr3* in the shRNA transgenics lies at a critical threshold. If a minimal threshold of XLR3 expression is met, MSCI succeeds and meiocyte loss is averted; but if XLR3 levels are just below the threshold, MSCI is abrogated and the spermatocytes undergo apoptosis. Spermatocytes from shRNA-*Xlr3* transgenic mice reveal a range of phenotypes that model these predictions: from cells with XY body MSCI marks well below the levels of wild type, likely contributing to the increased apoptotic figures observed; to cells with normal MSCI mark intensity that presumably go on to produce viable sperm (Fig. 4H-J, Fig. 7). It should be noted that the fact that *Xlr3* genes are themselves subject to MSCI lends an additional dynamic to the phenotypic variance seen in the knockdown.

The apparent dosage sensitivity of *Xlr* family members may also offer an explanation for why one *Xlr3* paralog, *Xlr3b,* is imprinted [28]. In females, the paternal allele of *Xlr3b* is repressed in both fetal and adult tissues, including ovary. Imprinting of the paternal copy of *Xlr3b* may serve to lower overall *Xlr3* expression in oocytes below the threshold where it might induce chromatin silencing inappropriately. Aneuploid mouse models exhibiting unpaired chromatin in females and/or unpaired chromatin apart from the sex chromosomes in males may shed light on this hypothesis.

## Materials and methods

### Ethics statement

All mouse procedures were approved by the Institutional Animal Care and Use Committee of the University of Connecticut.

### Sequence homology and alignments

Sequences were downloaded from the PFAM database (Family: Cor1 [PF04803]). Percent identity matrices (PIMs) were generated using ClustalW. Protein domain alignments and corresponding graphics were generated using NCBI’s constraint-based multiple alignment tool (COBALT) [60]. The *Xlr3/4/5* locus map was drawn to scale based on the UCSC genome browser view of identified regions (mm10).

### Animal husbandry & transgenesis

Mice were housed under climate-controlled conditions with a 12-h light/dark cycle and provided standard food and water ad libitum. A floxed stop containing, short hairpin RNA (shRNA) complementary to all *Xlr3* transcripts was cloned into the ROSA-PAS gene targeting vector [61]. This vector was electroporated into C57BL/6J (RRID:IMSR 000664) mouse embryonic stem cells (mESCs) and targeted clones were screened by long range PCR and Southern analysis. Two independent clones (B2 and B11) were microinjected into B6(Cg)-Tyrc-2J/J (C57BL/6J albino) host embryos (RRID:IMSR JAX:000058). Germline transmission was achieved through multiple male chimaeras from clone B2, and a stable line was bred through three stable male founders by backcrossing with C57BL/6J wild-type female mice. All experiments were performed on the 6th to 13th generation, male and female offspring (Fig. S1). The C57BL/6J-Gt(ROSA)26Sor<tm1(shRNA:Xlr3)Lgr>/LgrJ mouse line will be available for distribution from The Jackson Laboratory as stock #034383. As described in Fig. S1, B6.Cg-Tg(Pgk1-flpo)10Sykr/J (RRID:IMSR 011065) and FVB-Tg(Ddx4-cre)1Dcas/J (RRID:IMSR 006954) were used in this study, along with C3H/HeJ (RRID:IMSR 000659) mice.

### Tissue collection, testis weighing, and sperm counting

Tissues collected from euthanized mice were fixed as described in the following sections or snap frozen in liquid nitrogen and stored at −80°C. Testis weights were taken from fresh tissue before freezing or fixation. Sperm counting was performed as described by Handel (Handel and O’Brien, https://phenome.jax.org/projects/Handel1/protocol).

### PCR genotyping

DNA was extracted from ear punches by Proteinase K digestion. All primers used in this study are listed in Supplementary Table 1 with respective annealing temperatures. To amplify the shRNA-*Xlr3* template, PrimeSTAR GXL (Clontech) enzyme system was used as per manufacturer instructions (cycling conditions: 98°C for 10s, 60°C for 15s and 68°C for 10s/kB – products 3-4kB for 30s and <1 kB for 10s, for 35 cycles). Products were visualized on 1% agarose Tris-acetate-EDTA gels. All other polymerase chain reactions (PCR) were performed using the GoTaq (Promega) enzyme system as per manufacturer instructions (95°C for 2 min, then 95°C for 30s, annealing as specified for 30s, 72°C for 1 min repeated for 35 cycles, then 72°C for 5 min) and products were visualized on 1-2% agarose and Tris-borate-EDTA gels.

### Xlr3 antibody generation

The first 18 residues common to both XLR3A and XLR3B, and one amino acid difference from XLR3C, were used for peptide synthesis in rabbit immunization by Biosynthesis Inc. Following initial immunization, two rabbits received five biweekly boosters. Affinity purification of total IgG sera was performed using an AminoLink resin column (Thermo) and purified antibodies were complemented with pre-immune sera.

### Western blotting

Whole testis lysate was made by homogenization in RIPA buffer (50mM Tris-HCl pH 7.4, 150mM NaCl, 1% Triton X-100, 0.5% sodium deoxycholate, 0.1% SDS, 1mM EDTA, and 1mM PMSF), incubation on ice for 30 minutes, and centrifugation. Nuclear and cytoplasmic testis fractions were prepared from fresh tissue using a dounce homogenizer in Harvest Buffer (10mM HEPES pH 7.9, 25mM KCl, 2M sucrose, 1mM EDTA, 0.5mM spermidine, 1x protease inhibitor, 10% glycerol). Protein lysates were resolved by 12% SDS-PAGE and transferred to PVDF membrane (GE Healthcare Amersham^TM^) using a Trans-Blot^®^ SD Semi-Dry Transfer Cell (BioRad). The membrane was blocked with 5% milk in Tris-buffered saline (TBS), incubated in primary antibody overnight, followed by secondary antibody for 45 min. Signal was detected by chemiluminescence (Western Lightning, Perkin Elmer) on autoradiography film. Antibodies used in this study are listed in Supplementary Table 2.

### Tissue fixation, sectioning, and staining

Testes were dissected from adult mice and fixed as described previously [62]. Paraffin embedding was performed by the University of Connecticut Veterinary and Medical Diagnostic Laboratory. Sections were sliced to 6 μm on a rotary microtome for slide mounting. Hematoxylin and eosin (H&E) regressive staining was performed with Modified Harris Hematoxylin and Eosin Y 1% stock solution (Ricca Chemical Company) and used following manufacturer’s instructions. Apoptosis detection was performed using the DeadEnd™ Colorimetric Apoptosis Detection System (Promega) following the manufacturer’s instructions for fixed tissue slides. Imaging was performed on a light field microscope at 10x magnification.

### Spermatocyte chromosome spreads and immunocytochemistry

Spermatocyte spreads were made as previously described [63] with several modifications. Testes were dissected from 13.5-18.5 dpp mice, detunicated, and free tubules were suspended in 1x phosphate buffered saline (PBS). Tubules were broken up and cells were released by needle disruption. A single cell suspension was made by putting the tubule mixture through a 70 μm filter. Cells were placed in hypotonic solution of 2x HEB, centrifuged, and resuspended in fixative (1% PFA, 0.15% Triton X-100, and 13 mM DTT). Cells were collected by centrifugation and resuspended in 3.4 M sucrose, then applied to slides under 50-75% humidity. Slides remained in humidity for approximately 2 hours to allow cells to spread, then were washed in 1x PBS, and air dried.

Spermatocyte spreads were permeabilized with 0.5% Triton X-100 and blocked with 10% goat serum. Slides were incubated with diluted primary antibody at 4°C overnight and subsequently with diluted secondary antibody at 4°C for 30 minutes. Antibodies used in this study are listed in Supplementary Table 2. 4’6’-diamidino-2-phenylindole (DAPI) diluted 1:5 in Vectashield mounting media (Vector Labs) was used as a counterstain. Imaging was performed using an oil-immersion 100x objective of the Olympus IX 71 (DeltaVision) fluorescent microscope with DAPI, FITC, and TRITC channels. Images were captured and deconvolved using the softWoRx software (Applied Precision, LLC). Exposure times were set for particular antibody combinations and maintained for all slides imaged. Quantification of signal was accomplished by outlining cells and signals to take area and intensity measurements using ImageJ (Schindelin et al., 2015). Corrected total fluorescence (CTF) was calculated using the following formula:

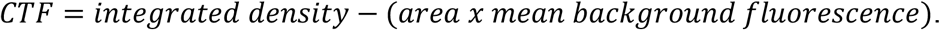

### RNA isolation and quantitative reverse transcription PCR (qRT-PCR)

RNA was extracted from frozen tissue using the NucleoSpin RNA kit (Machery-Nagel) as described by the manufacturer. cDNA was synthesized from total RNA using the QSCRIPT cDNA Supermix (Quanta Biosciences) and qRT-PCR was carried out in triplicate using the iTAQ Universal SYBR Green SuperMix (BioRad). Primers for these reactions are listed in Supplementary Table 1 and all reactions were carried out under the following cycling conditions: initial denaturation at 95°C for 3 mins, 40 cycles (95°C for 5 sec 58°C for 30 sec), a final extension of 65°C for 5 sec. CT and melting curve results were calculated by the CFX Manager 3.1 software (BioRad). Results were normalized to *β-actin* or *Gapdh* using the ΔΔCt method [64].

### RNA fluorescence *in situ* hybridization (FISH)

RNA FISH was performed as described by [65]. Probes were generated using primers listed in Supplementary Table 1 from the G135P65476A4 BAC as described in [8] and were labeled using the DIG-Nick Translation Mix (Roche) as per manufacturer’s instructions. Following RNA FISH, slides were fixed in 4% paraformaldehyde and processed for IF as above.

### Graphical and statistical data presentation

All boxplot and bar graphs were produced using Rstudio (Rstudio Team, 2020). In the case of bar graphs, all error bars represent the mean ± confidence interval (CI). In the case of boxplots, unfilled dots represent outliers and colored dots represent individual observations. Normal data distribution was tested using the Kolmogorov-Smirnov test. To determine statistical significance, if data were normally distributed, a T-test was applied; if data were not normally distributed, a Wilcox Test was applied. All sample sizes are indicated with the relevant tests.

## Acknowledgements

We thank the members of the M.O’Neill and R.O’Neill laboratories for their discussion and helpful comments. Thank you to R. O’Neill for critical evaluation of the manuscript, N. Pauloski and C. McCann with microscopy. Thank you to Dr. Satoshi Namekawa for the gift of the anti-BRCA1 antibody used in this study.

---

**Supplementary Figure 1.**
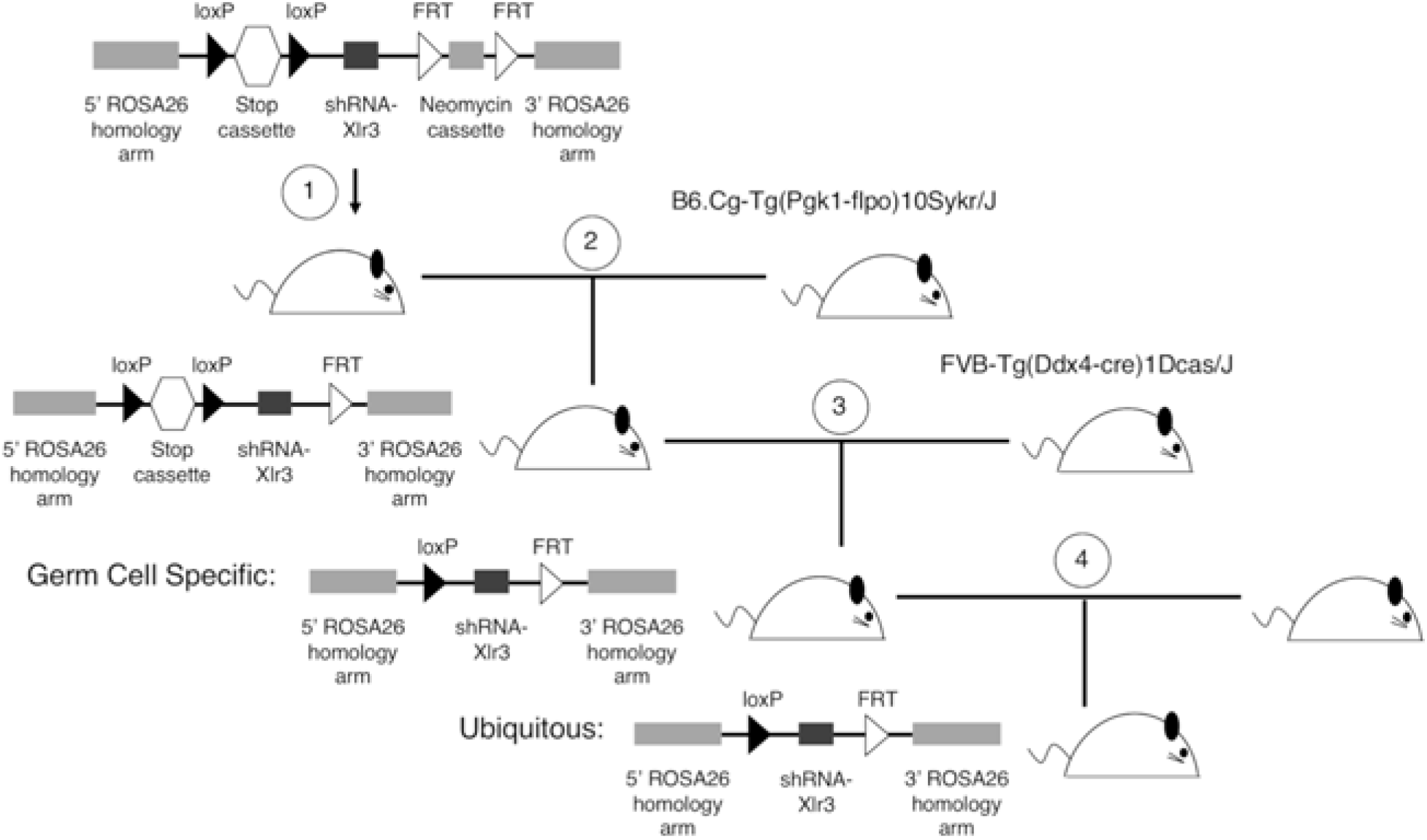
Breeding schematic of shRNA-*X/r3* transgenic mouse. 1) The linearized vector was electroporated into cells and incorporated into the *ROSA26* locus on chromosome 6 by homologous recombination. Chimeric mice were produced by blastocyst incorporation. 2) Mice with full-length construct were crossed to mice expressing flippase to recombine the *FRT* sites surrounding the neomycin cassette. 3) Mice with shortened construct were crossed to *Ddx4-Vasa-Cre* mice expressing Cre recombinase in germ cells. The stop cassette was excised and shRNA-*X/r3* was expressed in a tissue-specific manner. 4) Ubiquitously expressing shRNA-*X/r3* mice were generated by breeding mice with the active construct in the germ line.

**Supplementary Figure 2.**
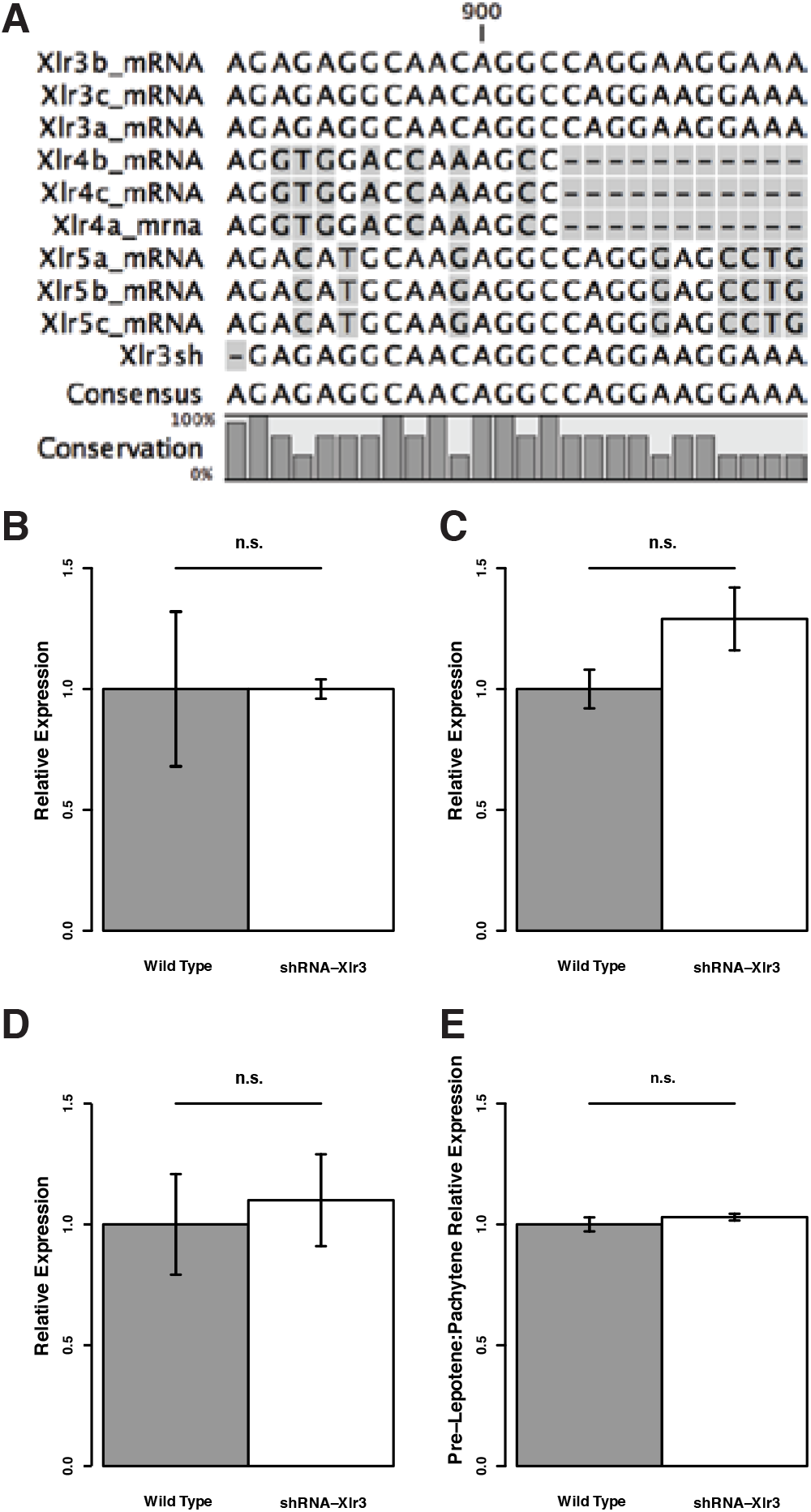
Off-target effects as a result of shRNA-*X/r3* activation were not observed. **A)** Alignment and conservation of *X/r3/4/5* mRNA sequences at the site of shRNA targeting generated by CLC Sequence Viewer 7 (Qiagen). **B)** *X/r4* (T-test: t=-0.0217, df=11.953, p-value=0.983), **C)** *X/r5* (T-test: t=-2.7428, df=10.204, p-value=0.7893), **D)** *Oas1b* (T-test: t=-0.68427, df=11, p-value=0.508), **E)** and *Sox9* (T-test: t=0.44358, df=9.5631, p-value=0.6672) transcription levels as measured by qRT-PCR are not significantly (n.s) different between wild type and shRNA-*X/r3* testis total cDNA. Error bars represent mean C.I.

**Supplementary Table 1.**
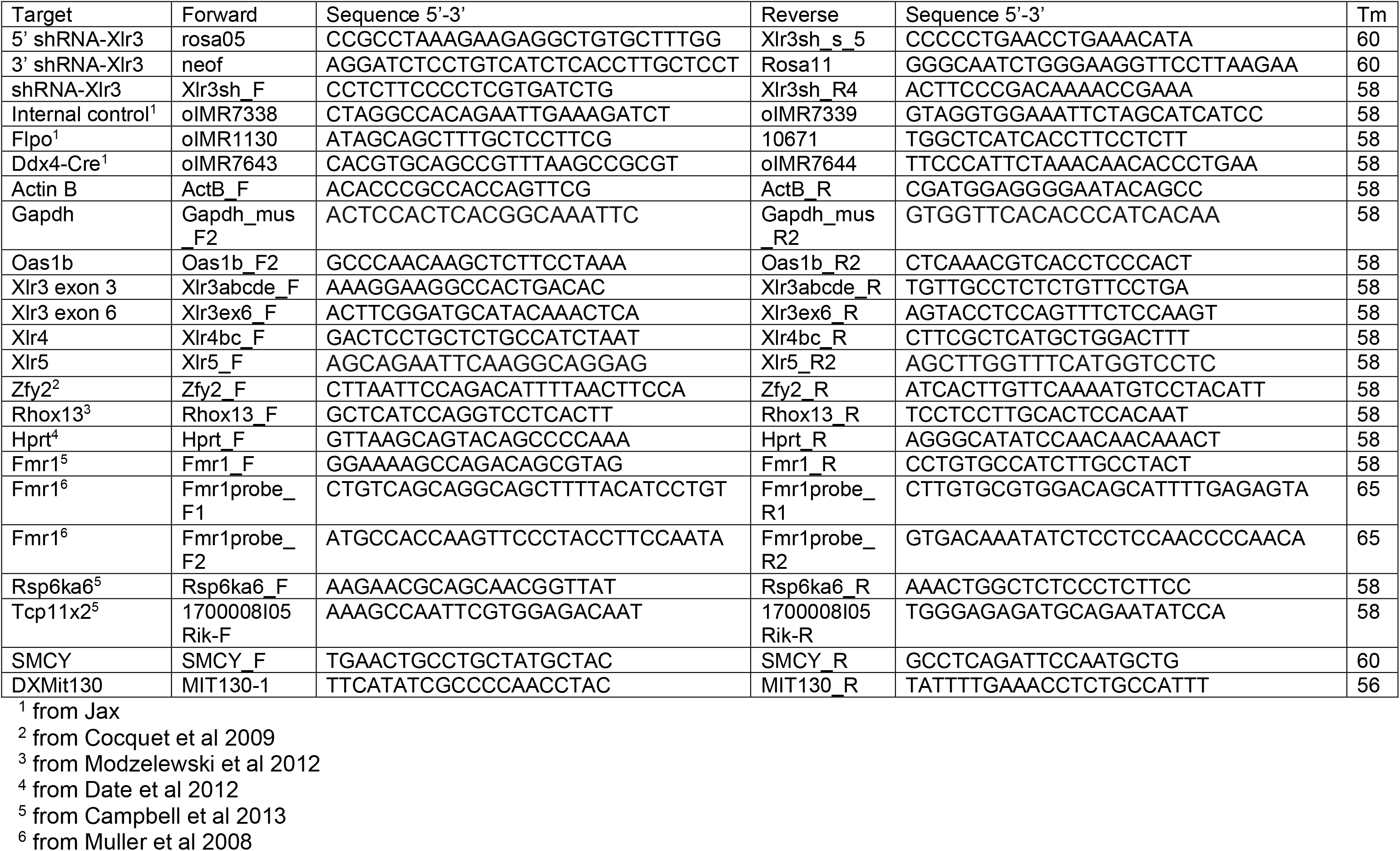
Primers used in this study

**Supplementary Table 2.** Antibodies used in this study

| Antibody | Concentration | Supplier | Catalog #/RRID |
| --- | --- | --- | --- |
| Rabbit polyclonal anti-Xlr3 | ICC 1:20<br>Western 1:2,000 | Custom generated by Biosynthesis Inc to peptide MSSRK RKATSTAGRHSRM | N/A |
| Mouse monoclonal anti-Sycp3 | ICC 1:200 | Santa Cruz | sc-74569<br>RRID:AB_2197353 |
| Rabbit polyclonal anti-Sycp3 | ICC 1:100 | Thermo Fisher | PA1-16764<br>RRID:AB_568727 |
| Mouse monoclonal anti-gH2aX (ser139) | ICC 1:200 | Thermo Fisher | 14-9865-82<br>RRID:AB_2573048 |
| Rabbit polyclonal anti-Sumo1 | ICC 1:200 | Thermo Fisher | PA5-17352<br>RRID:AB_10977036 |
| Rabbit polyclonal anti-H3K9me3 | ICC 1:180 | Thermo Fisher | 49-1008<br>RRID:AB_2533859 |
| Rabbit polyclonal anti-α-tubulin | Western 1:40,000 | Abcam | Ab15246<br>RRID:AB_301787 |
| anti-rabbit IgG HRP | Western 1:2,000 | Abcam | Ab6721<br>RRID:AB_955447 |
| anti-mouse IgG HRP | Western 1:100 | Abcam | Ab6728<br>RRID:AB_955440 |
| Goat anti-Mouse Alex Fluor® 568 | ICC 1:200 | Invitrogen | A11004<br>RRID:AB_2534072 |
| Goat anti-Rabbit Alex Fluor® 488 | ICC 1:200<br>RNA FISH 1:200 | Invitrogen | A27034<br>RRID:AB_2536097 |
| Anti-DIG Fluorescein | RNA FISH 1:200 | Roche | 11207741910<br>RRID:AB_514498 |
| Rabbit anti FITC | RNA FISH 1:200 | Invitrogen | 71-1900<br>RRID:AB_2533978 |

